# Long-term aberrations to cerebellar endocannabinoids induced by early-life stress

**DOI:** 10.1101/830901

**Authors:** Alexandra B. Moussa-Tooks, Eric Larson, Alex F. Gimeno, Emma Leishman, Lisa A. Bartolomeo, Heather Bradshaw, John T. Green, Brian F. O’Donnell, Ken Mackie, William P. Hetrick

## Abstract

Studies of early-life stress traditionally focus on glucocorticoid signaling as a modulator of neurodevelopmental risk, but emerging evidence points to the role of the endocannabinoid system in long-term stress-induced neural remodeling. Existing studies on stress-induced endocannabinoid dysregulation have focused on changes to cerebrum that are temporally proximal to stressors, but little is known about temporally distal effects, especially in cerebellum, which is vulnerable to early developmental stress and is dense with cannabinoid receptors. Further, sex-specific effects of stress on cerebellar endocannabinoid tone are understudied. Following a naturalistic rodent model of early-life stress, limited bedding at postnatal days 2-9, adult (postnatal day 70) cerebellar and hippocampal endocannabinoids and related lipids and mRNA were assessed, and behavioral performance was evaluated. Regional and sex-specific effects were present at baseline and following early-life stress. Limited bedding impaired peripherally-measured basal corticosterone in adult males only. In the CNS, early-life stress (1) decreased 2-arachidonoyl glycerol and arachidonic acid in the cerebellar deep nuclei in males only; (2) decreased 2-arachidonoyl glycerol in females only in cerebellar Crus I; and (3) increased dorsal hippocampus prostaglandins in males only. Transcriptomics for cerebellar interpositus nucleus revealed substantial sex effects, with minimal effects of stress. Stress did impair novel object recognition in both sexes and social preference in females. Taken together, the cerebellar endocannabinoids system exhibits robust sex-specific differences, malleable through early-life stress and perhaps also contributing to sexual differentiation of the brain. The current study may foster future research into stress as a risk factor for cerebellar-related dysfunctions.

## Introduction

In humans, poor prenatal and early postnatal care is a major developmental stressor commonly found in low socioeconomic groups ^1,2^. Such stress can affect neurodevelopment during critical periods of early life, greatly impacting both cognitive and behavioral outcomes in adulthood ^3-8^. Many studies have examined the effects of stress on the cerebrum, such as the hippocampus, and found both neural and behavioral aberrations ^9^. However, compelling evidence shows that the cerebellum, which is bidirectionally interconnected with vast regions of the forebrain ^10-14^, is also profoundly affected by stress ^15,16^.

### The Stress Response

Evidence suggests that developmental impairments in humans and rats are mediated by poor regulation of homeostatic systems as a function of early-life stress ^17,18^. One such homeostatic mechanism is the stress response. Stress typically results in hypothalamic release of corticotropin releasing hormone (CRH), which stimulates the pituitary gland to secrete adrenocorticotropic hormone (ACTH) and, in turn, prompts the adrenal cortex to release cortisol (or the rat homologue corticosterone; CORT). CORT enters the bloodstream and acts on various nervous system targets, which provides negative feedback on the hypothalamic-pituitary-adrenal (HPA) axis, including the hypothalamus. CORT binds to glucocorticoid receptors in hypothalamic periventricular nucleus (PVN) neurons, a process that ultimately suppresses the HPA axis by inhibiting the continued release of glutamate. The endocannabinoid system plays an important role in this regulation ^19,20^.

### Endocannabinoid Regulation of Stress

Recent studies indicate that stress may increase risk for the varied detrimental effects of exogenous cannabinoids like Δ^9^-tetrahydrocannabinol (THC) ^21^. Accordingly, it seems imperative to understand the vulnerability that stress imparts on neural systems. Heavily integrated into the stress response system is the endogenous cannabinoid (endocannabinoid) system. Once CORT has activated post-synaptic glucocorticoid receptors, those receptors signal the production of endocannabinoids, such as 2-arachidonoyl glycerol (2-AG) and *N*-arachidonoyl ethanolamine (anandamide; AEA). Endocannabinoids primarily act in a retrograde manner, signaling at presynaptic receptors to modulate synaptic transmission and the stress response ^20^. CB_1_ cannabinoid receptors (CB1Rs) are present at high levels in the prefrontal cortex, hippocampus, and cerebellum, modulating transient and long-lasting forms of synaptic plasticity ^22,23^ and the stress response ^24^. CB1Rs are more abundant in early life as the endocannabinoid system undergoes significant remodeling and refinement, suggesting that this system is particularly vulnerable during this time of plasticity ^25^.

Just as there are widely known sex differences in the stress-response system, including basal CORT levels, stress responsivity, and glucocorticoid receptors ^26^, there is emerging evidence of sex differences in the endocannabinoid system. For example, there appear to be more CB1R protein in males than females, though females tend to show higher CB1R G-protein activation from endocannabinoids, suggesting more efficient coupling of CB1R in females ^25^. These sex differences in the cannabinoid system may be important for key sex differences observed in behavior following stressors. Early-life stress affects the endocannabinoid system by (a) decreasing CB1Rs in adulthood in males ^27^ across various cortical regions as concluded through [^3^H]CP55,940 binding and mRNA expression, though findings in females have been mixed ^28^, and (b) increasing gene expression for endocannabinoid degradation enzymes ^29^, though surprisingly these effects have not been studied in the cerebellum. Although endocannabinoids have not been thoroughly investigated in the cerebellum, males have been shown to exhibit increased cerebellar 2-AG compared to females ^30^.

### Neural Vulnerability to Stress

Perhaps unsurprisingly, neural regions highly sensitive to stress are also dense with endocannabinoid receptors. The hippocampus, which has received substantial attention in this field, is well known for its association with the formation of declarative and spatial memories. Hippocampal memory formation appears to be highly susceptible to stress, likely due to its numerous inhibitory connections with the HPA axis ^31,32^. Moreover, the hippocampus manifests one of the highest affinities for cortisol binding ^33,34^. Human studies indicate that early-life stress is associated with decreased hippocampal volume and activation ^5,35,36^. Similarly, animal models of stress exhibit dendritic atrophy and impaired stress-hormone receptor binding and expression ^37^ as well as delayed milestones for dentate gyrus development ^38^ in the hippocampal formation following limited bedding. The hippocampus’s role in memory formation and linkage with the HPA axis may explain the array of impairments common to early-life stress and the development of psychopathology ^39^.

Another region dense in CB1Rs is the cerebellum, which has traditionally been understudied in the stress literature. What little evidence exists regarding the sensitivity of this region to stress is compelling. The cerebellum undergoes cellular differentiation even into 2 years of age in humans or 3 weeks in rats, with continued proliferation, development, and reorganization occurring well into adolescence and young adulthood, and is likely vulnerable to early-life experiences during this time ^40^.

Research in human populations has indicated that developmental insults, including preterm birth ^41,42^ and early-life maltreatment ^43-45^, are related to impairments in cerebellar development, findings which have been corroborated in animal models ^15^. For example, Llorente and colleagues ^46^ found that early-life stress induced cerebellar neuronal degeneration in adult rats, most prominently in males, that experienced this stressor as pups ^46^. Wilber and colleagues ^47^ revealed an increased number of glucocorticoid receptors within the cerebellar interpositus nucleus after a maternal deprivation early-life stress paradigm that was correlated with a marked behavioral deficit in delay eyeblink conditioning (EBC), a cerebellar-dependent task ^47-49^. Thus, early-life stress appears to meaningfully impact the cerebellum, its development, and its associated learning processes.

In addition to the interpositus nucleus, EBC requires cerebellar cortex near the base of the primary fissure. Cerebellar cortex is critical for other cognitive functions, creating closed-loop circuits with prefrontal cortex and other cerebral areas that may allow for cerebellum to modulate higher-order functions ^50,51^. For example, Crus I has been linked to cognitive flexibility ^52^, perceptual decision-making ^53^, and social behaviors ^54^.

Chronic stress across the life span remodels the hippocampal endocannabinoid system, by increasing 2-AG, which has been linked to increased CORT ^20^. However, despite the abundance of cannabinoid receptors in the cerebellum and clear evidence of its susceptibility to stress, it remains unknown how early-life stress impacts cerebellar endocannabinoids. Further, it has been suggested that neurodevelopmental changes following early-life stress may not be detectable until adulthood ^55,56^, which brings into focus the importance of studying long-term effects of early-life stress. The current study, for the first time, concurrently examined the effects of early-life stress on endocannabinoids in the hippocampus and cerebellum, both of which are nodes of endocannabinoid signaling and vulnerable to early-life stress.

Here, a naturalistic rat model of maternal stress, limited bedding, was used to test the hypothesis that early-life stress results in long-term changes in the cerebellar endocannabinoid system. Given the amplification of the stress response during this critical early-life period of cerebellar and endocannabinoid system development, as measured by CORT, it was predicted that early-life stress would down-regulate endocannabinoids in adulthood as measured by lipid analysis. Similarly, it was expected that cerebellar cortex and dorsal hippocampus would both show changes in key endocannabinoids, including increased 2-AG and decreased AEA, as has been shown in hippocampal studies of early-life stress, with accompanying changes to cannabinoid-related mRNA. At baseline, females were expected to have lower endocannabinoid and higher CORT levels compared to normally reared males, following previously reported sex differences. Stressed animals were also expected to exhibit impairments in key behavioral processes including recognition memory and social preference, based on previous studies ^57^. Moreover, these behavioral differences were not expected to be due to an anxiety phenotype (assessed via elevated plus maze) or gross cerebellar dysfunction (assessed via rotarod). Overall, these predictions form a direct test of the model that early-life stress induces vulnerability of a portion of the cerebellar-endocannabinoid network, with implications for long-term, aberrant behavioral outcomes.

## Methods

All experimental protocols involving rats were approved by the Indiana University Bloomington IACUC.

### Animals

Timed-pregnant Long-Evans rats (Envigo, Indianapolis, IN) arrived at gestational day 14. Dams were individually housed in polypropylene cages (26.67cm×48.26cm×20.32cm) in a 12:12-hour light-dark cycle (6:00 lights on, 18:00 lights off) and temperature-controlled (22.8°C) vivarium. Food and water were provided ad libitum. Bedding was changed once per week. Rats were checked every 12 hours surrounding the expected date of birth. Day of birth was designated P0. At P2, animals were randomly cross-fostered, and cages were sex-balanced.

### Limited Bedding Stressor

At P2, all cages were changed. Half of the cages were randomly selected to undergo the limited bedding manipulation. Limited bedding cages contained a wire mesh insert (Plastic-coated aluminum mesh, 0.4cm×0.9cm, McNichols Co., Tampa, FL) that was fitted 2.5cm above the cage floor (c.f., ^57^). The mesh allowed the passage of excrement to the bedding material below the mesh. Additionally, limited bedding cages were given half of a paper towel square (13.97cm×27.94cm) for the dam to use as nesting material. Normal rearing cages were given a full paper towel square and standard access to bedding material. All cages were left undisturbed from P2-9. On P10, all animals were transferred to clean, standard caging. All animals were reared normally from this point forward. At P21 pups were weaned into treatment- and sex-matched cages of 3-4 animals.

### Plasma Corticosterone Quantification

To assay acute and long-term changes in basal stress, plasma corticosterone levels were quantified using trunk blood. Animals were sacrificed at P8 (during the stressor) using rapid decapitation and P70 (adulthood) using isoflurane then rapid decapitation. After decapitation, trunk blood was collected in 1.5mL Eppendorf tubes with 1μL of liquid heparin (1 000 USP/mL) to prevent clotting and immediately frozen at -80°C. Samples were shipped to Cayman Chemicals for quantification via ELISA (sensitivity 30pg/mL; intra-assay variation 2%–18%).

### Endocannabinoid Quantification

Upon sacrifice at P70, neural tissue was harvested, frozen, and sectioned using a 1mm coronal sectioning block (BrainTree Scientific). Two-millimeter round punches were taken across three adjacent sections of the interpositus (IP) nucleus and Crus I of the cerebellum and dorsal hippocampus. Punches were flash frozen using liquid nitrogen and stored at -80C. To extract and partially purify lipids from these tissues, samples were processed by the Bradshaw Lab of Lipid Neuroscience at Indiana University Bloomington as previously described (c.f., ^58^).

26 lipids from the broader endocannabinoid lipidome, including AEA, its lipoamine structural analogs (e.g., *N*-palmitoyl ethanolamine), 2-AG, its 2-acyl glycerol structural analogs (e.g. 2-linoleoyl glycerol), associated free fatty acids, and prostaglandins were quantified in tissue extracts from a single hemisphere using high-performance liquid chromatography coupled with tandem mass spectrometry (HPLC/MS/MS). Lipid concentrations were normalized to sample mass prior to statistical analysis. Group comparisons were performed via one-way ANOVAs, correcting for multiple comparisons.

### mRNA Analysis

Two-millimeter round punches from the corresponding hemisphere used for lipid analysis were taken across three adjacent sections of the IP nucleus. Tissue samples were immediately homogenized in Trizol in preparation for mRNA isolation. Purification (QIAGEN RNeasy Plus Mini Kit) yielded 20μL of RNA, which was quantified using Nanodrop and assessed for integrity via an Agilent 2100 Bioanalyzer. Three samples were eliminated due to low RNA integrity (RIN<5). Samples were processed, run, and analyzed by the Indiana University School of Medicine Genomics Core Facility as described below.

A library was prepared using KAPA mRNA Hyperprep Kit (KK8581). Sequencing was performed using Illumina HiSeq 4000 sequencing and deemed good quality (95% Q30, 76% Cluster Density, 367M reads/lane). Alignment and mapping also yielded acceptable quality (85.5% uniquely mapped reads, average 49.4% mapped onto gene). Differential expression analyses were performed using treatment and sex as primary factors and corrected for multiple comparisons.

### Behavioral Assessment

Pups were assessed on four behavioral tasks in young adulthood. At P46, all rats underwent 3 consecutive days of handling and familiarization to the testing room. Behavioral testing continued from P50-70. Thirty rats underwent Novel Object Recognition (NOR), then Rotarod. 44 rats underwent NOR, followed by Social Preference, then Elevated Plus Maze (see Fig. 5 for group sizes).

**Figure 1.**
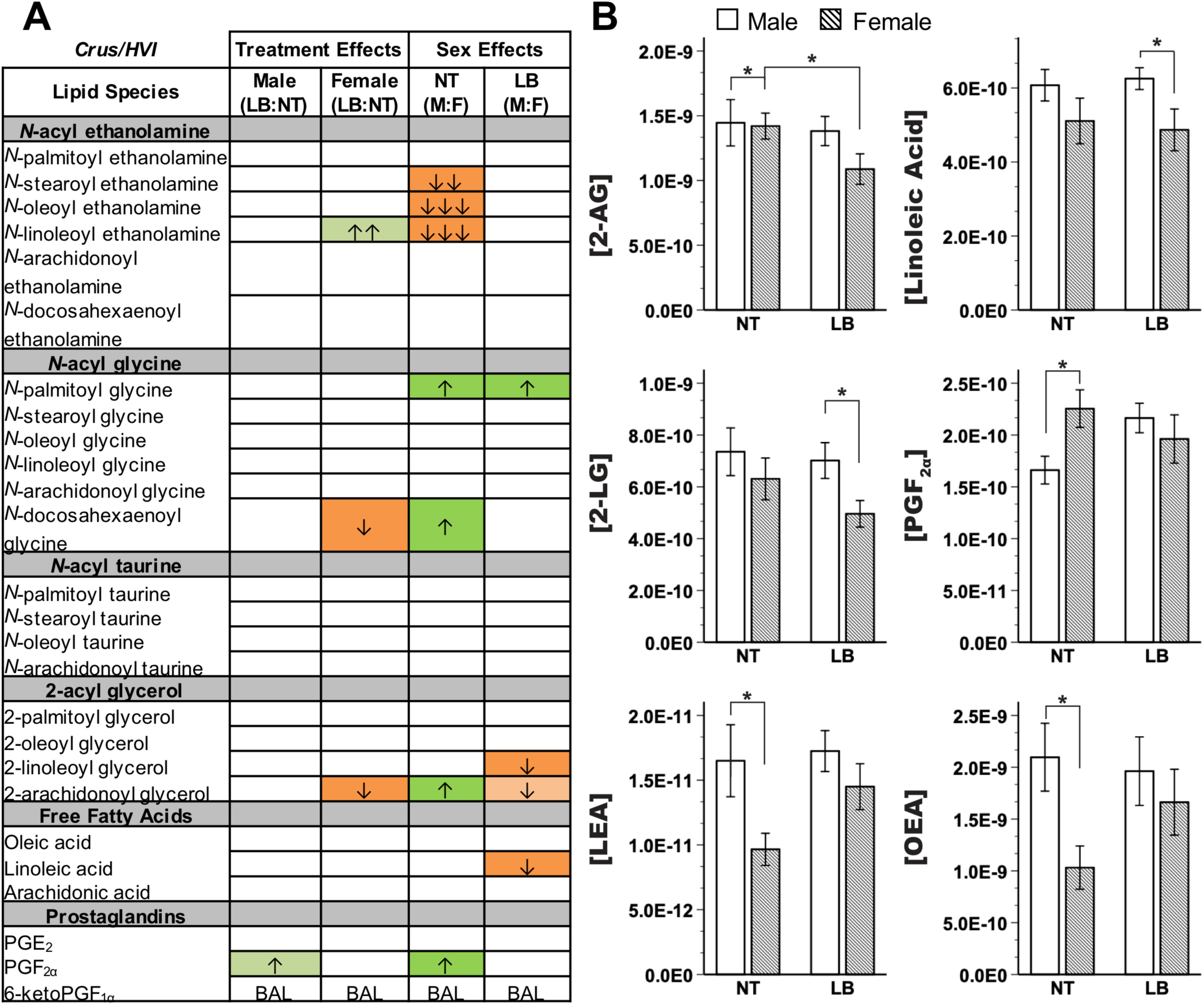
Endocannabinoid and related lipid quantification for the cerebellar Crus I region, showing (**A**) effects for all measured lipids. Columns represent treatment and sex-specific effects, showing changes to the limited bedding group in relation to the no treatment group (LB:NT) or changes in males in relation to females (M:F). Number of arrows represents fold-change: up-arrow (green) is increase, down-arrow (orange) is decrease; ↑=0-0.49-fold change, ↑↑=0.5-0.99-fold change, ↑↑↑=1-1.49-fold change. Color scheme represents significance, with full colors indicating significance at LSD-corrected *p*<0.05 and shaded colors indicating a trending significance. BAL=below analytical limits. (**B**) specific effects on lipids of interest. Y-axis indicates concentration of lipid of interest while x-axis represents limited bedding (LB) and no treatment (NT) groups. OEA=*N*-oleoyl ethanolamine, LEA=*N*-linoleoyl ethanolamine. *LSD-corrected *p*<0.05

**Figure 2.**
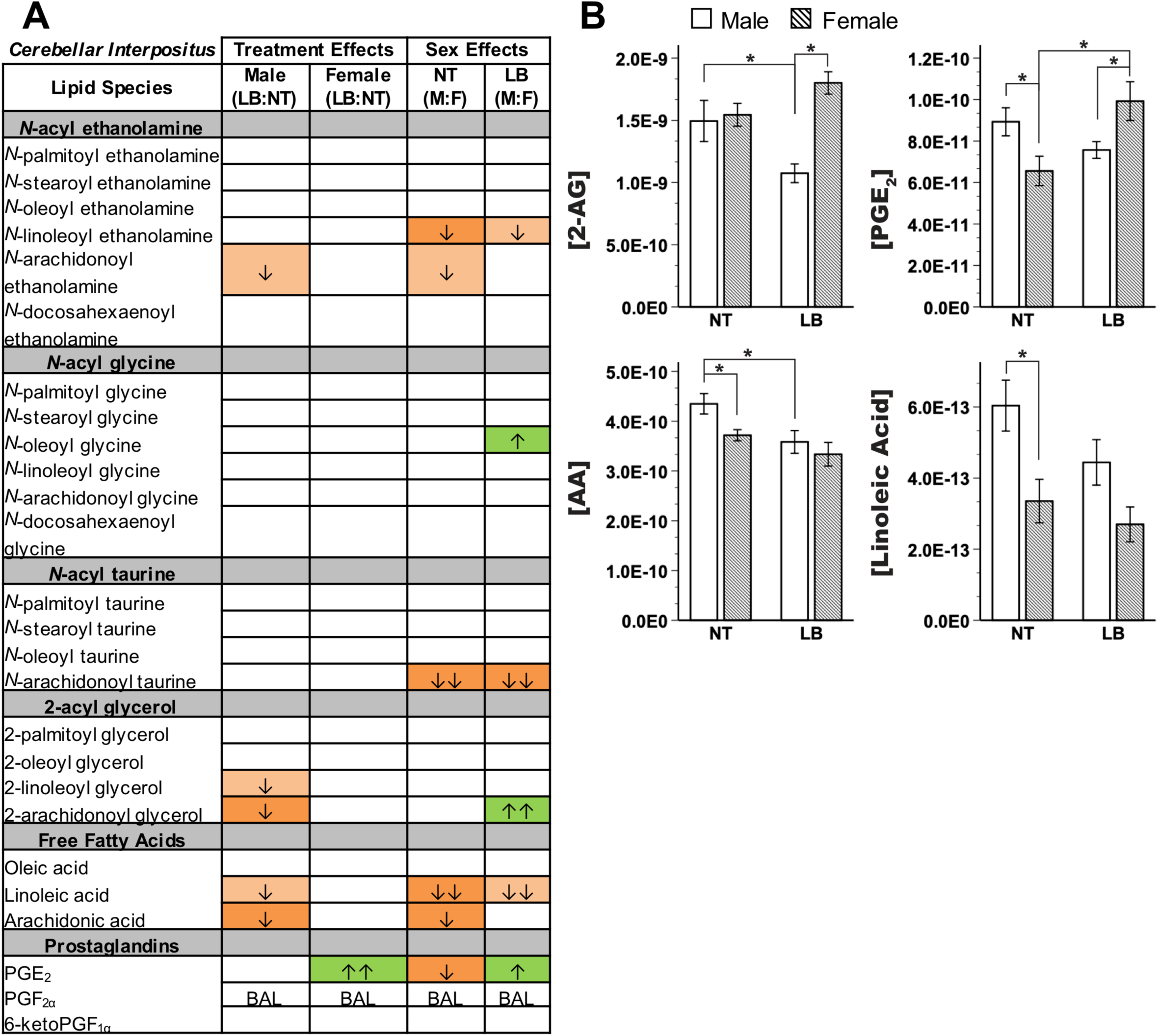
Endocannabinoid and related lipid quantification for the cerebellar interpositus (IP) region, showing (**A**) effects for all measured lipids. Columns represent treatment and sex-specific effects, showing changes to the limited bedding group in relation to the no treatment group (LB:NT) or changes in males in relation to females (M:F). Number of arrows represents fold-change: up-arrow (green) is increase, down-arrow (orange) is decrease; ↑=0-0.49-fold change, ↑↑=0.5-0.99-fold change. Color scheme represents significance, with full colors indicating significance at FDR corrected *p*<0.05 and shaded colors indicating a trending significance. BAL=below analytical limits. (**B**) specific effects on lipids of interest. Y-axis indicates concentration of lipid of interest while x-axis represents limited bedding (LB) and no treatment (NT) groups. *LSD-corrected *p*<0.05

**Figure 3.**
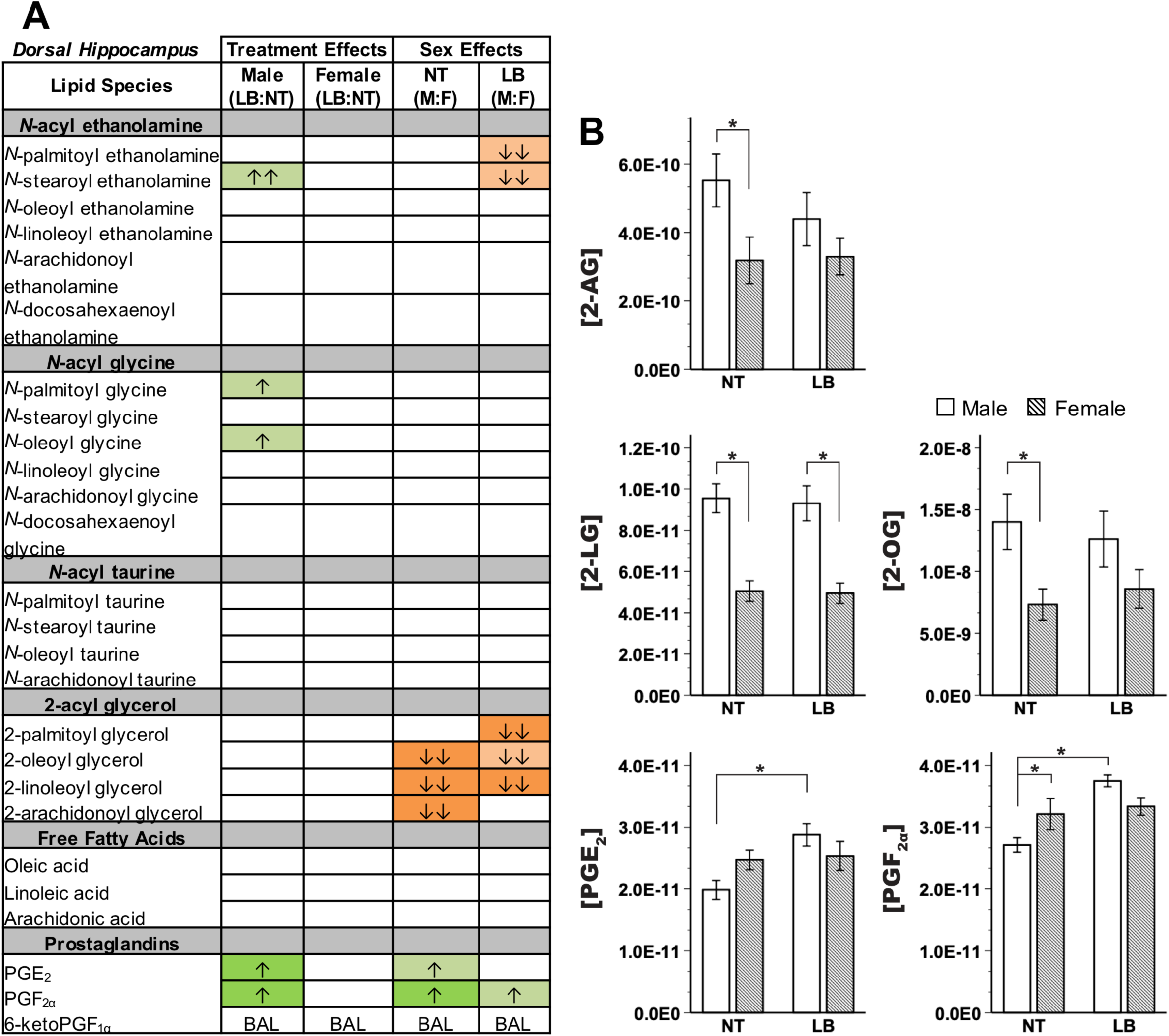
Endocannabinoid and related lipid quantification for the dorsal hippocampus region, showing (**A**) effects for all measured lipids. Columns represent treatment- and sex-specific effects, showing changes to the limited bedding group in relation to the no treatment group (LB:NT) or changes in males in relation to females (M:F). Number of arrows represents fold-change: up-arrow (green) is increase, down-arrow (orange) is decrease; ↑=0-0.49-fold change, ↑↑=0.5-0.99-fold change. Color scheme represents significance, with full colors indicating significance at LSD-corrected *p*<0.05 and shaded colors indicating a trending significance. BAL=below analytical limits. (**B**) Specific effects on lipids of interest. Y-axis indicates concentration of lipid of interest while x-axis represents limited bedding (LB) and no treatment (NT) groups. *LSD-corrected *p*<0.05

**Figure 4.**
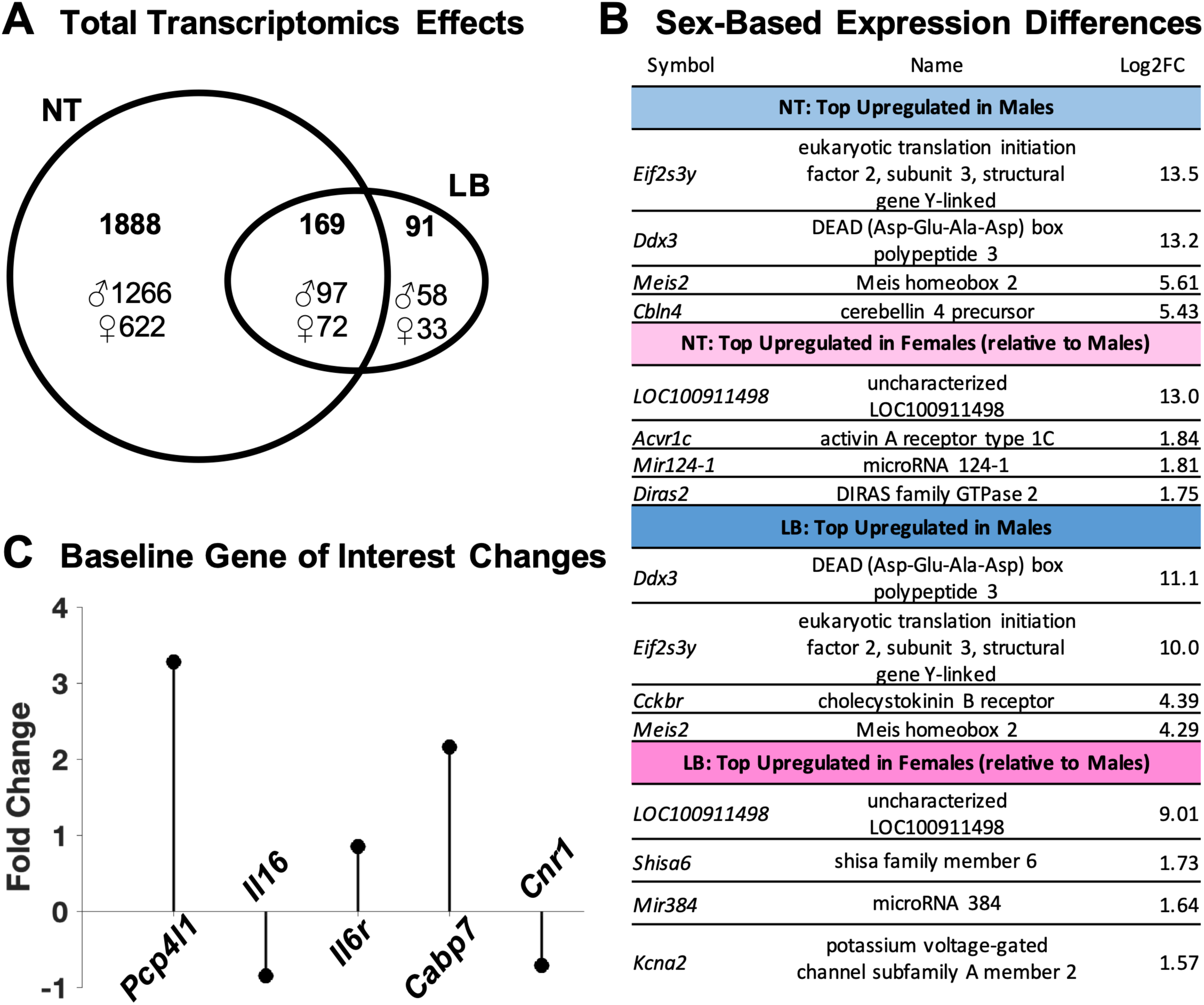
Summary of transcriptomics data for the cerebellar interpositus region. (**A**) Venn diagram depicting the number of genes significantly differing by sex within the no treatment (NT) or limited bedding (LB) groups and those impacted genes that are common between the two treatment groups. Top number in **bold** indicates the total number of genes changed, with ♂ and ♀ indicating number of genes upregulated in males and upregulated in females, respectively, as evidenced by log2 fold change data. (**B**) List of top upregulated genes in males and females within each treatment group based on the largest magnitude significant difference in expression, reported as log2 fold change (Log2FC). (**C**) Diagram depicting baseline significant expression differences (fold-change for five genes of interest, due to the relevance to the interpositus or cannabinoid system) between no treatment males and no treatment females. Positive values indicate upregulation in males while negative values indicate upregulation in females.

**Figure 5.**
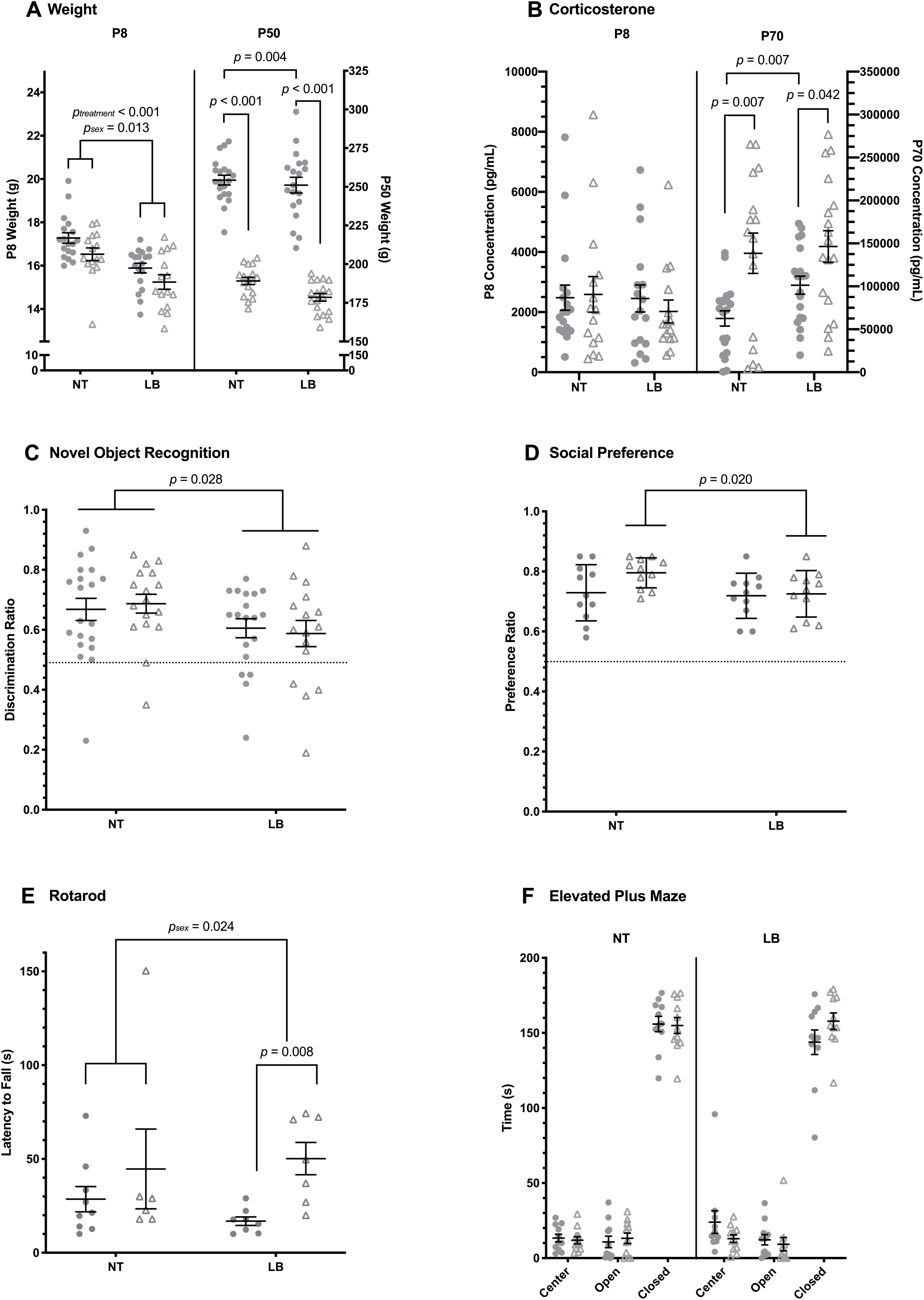
Behavioral data. (**A**) Weight across treatment groups (NT=no treatment; LB=limited bedding) at postnatal day 8 (P8, ⍰ =male [N=18 NT, 18 LB], Δ =female [N=15 NT, 15 LB]) and P50 (male N=20 NT, 19 LB; female N=17 NT, 18 LB) with individual points plotted for individual animals. (**B**) Corticosterone N=15 NT, 15 LB) and P70 (male N=20 NT, 19 LB; female N=16 NT, 18 LB). (**C**) Discrimination ratio from novel object recognition across treatment groups (NT=no treatment; LB=limited bedding) with individual points plotted for individual animals (male N=20 NT, 19 LB, female N=17 NT, 16 LB). Dotted line (y=0.5) indicates no preference, with animals above showing preference for the novel object and animals below showing preference for the familiar (N=11/group/sex). (**D**) Preference ratio from social preference test. Dotted line (y=0.5) indicates no preference, with animals above showing preference for the novel partner and animals below showing preference for the novel object. (**E**) Latency to fall during rotarod task (male N=9 NT, 8 LB; female N=6 NT, 7 LB). (**F**) Time spent in open, closed, and center arms of elevated plus maze (N=11/group/sex).

#### Novel Object Recognition (NOR)

Following handling, rats were familiarized to the NOR apparatus (15 minutes daily for 3 days) which included a clear Plexiglas box (50×50cm base, 40cm height; open top). During the testing day, rats were first given 10 minutes under red light to explore two identical objects located in adjacent corners of the testing apparatus. Rats were then returned to their home cage in the housing colony for a one-hour delay, after which they were placed back into the apparatus, under red light, for the test trial. One object from previously (“familiar”) and one novel object were located in adjacent corners of the box. Rats were given 5 minutes to explore. Blinded raters scored the first 3 minutes of the test trial for time spent interacting with the familiar and novel objects. A discrimination ratio was calculated (time spent with novel object/time spent with novel and familiar object) to quantify recognition memory.

#### Social Preference

Rats were exposed to a novel sex- and weight-matched rat restrained within a wire mesh box (11×11×19cm) and novel object covered by a wire mesh box in adjacent corners of the same apparatus as used for NOR (clear Plexiglas box) and in the same testing room to which they were previously familiarized. Blinded raters scored 3 minutes of interaction for time spent engaging with the rat (i.e., social) or object (i.e., non-social). A social preference ratio was calculated (time spent with novel rat/time spent with novel object and rat).

#### Rotarod

Following 3 days of familiarization to the apparatus (IITC Life Sciences, Roto-Rod Series 8), rats were placed on the center rod and allowed to move freely as the velocity increased from a starting point of 4 RPMs by 7.2 RPMs. Latency to fall was measured for each rat by a sensor within the apparatus.

#### Elevated Plus Maze (EPM)

The apparatus consisted of two open and two closed, elevated arms (each 30cm long). Rats were placed in the center of the platform facing an open arm and allowed 3 minutes to explore. Raters blinded to experimental group scored time spent in open arms, closed arms, and the center.

### Statistical Analyses

Using SPSS (25.0, IBM Corporation), 2-Way ANOVAs were used to assess main and interaction effects of treatment and sex on corticosterone, lipid, and behavioral outcomes. Following, correlation analyses were performed between variables yielding group treatment or sex differences to determine the relationship between biochemical features and behavioral outcomes.

## Results

### Weight

At P8, significant main effects of sex (F(1,62)=6.53, *p*=0.013, ηp^2^=0.095) and treatment (F(1,62)=24.68, *p*<0.01, ηp^2^=0.996) were observed, but no interaction effects were found (Fig. 5A). A significant main effect of sex (F(1,70)=384.49, *p*<0.01, ηp^2^=0.846) and trending main effect of treatment (F(1,70)=3.86, *p*=0.053, ηp^2^=0.052) were observed in adulthood (P50), with females weighing less than males, as expected, and limited bedding rats weighing less than normally reared rats, driven by the female group (t(33)=3.12, *p*=0.004).

### Plasma Corticosterone Quantification and Analysis

Consistent with the literature, a main effect of sex (F(1,69)=15.094, *p*<0.001, ηp^2^=0.179) was observed at P70, with females exhibiting higher CORT levels compared to males (Fig. 5B). At P70, males exhibited a significant treatment effect on systemic CORT such that limited bedding males had increased CORT concentrations compared to normally-reared males (t(37)=-2.836, *p*=0.007, d=0.907). No sex or treatment effects were observed at P8. Rats had significantly higher concentrations of systemic CORT at P70 than P8 (F(1,130)=176.566, *p*<0.001, ηp^2^=0.576).

### Endocannabinoid Quantification

Despite standardized tissue extraction volumes, sample mass showed significant sex differences in cerebellar regions (Supplementary Tables 2 and 3). Between-group effects were observed in Crus I (F(3,26)=5.74, *p*=0.004) with post-hoc, LSD-corrected comparisons revealing that normally reared females exhibited smaller mass compared to normally reared males (*p*=0.001) and in IP (F(3,26)=8.06, *p*=0.001), with normally reared females having smaller mass than normally reared males (*p*=0.02) and limited bedding females having smaller mass than limited bedding males (*p*=0.001) in the IP nucleus. Since all lipid concentrations were normalized to sample mass, these differences did not impact further analyses and interpretations of regional lipids.

Endocannabinoid levels in Crus I of the cerebellum showed a baseline sex difference (Figure 1, Supplementary Table 1): control females exhibited significant decreases in *N*-stearoyl ethanolamine, *N*-oleoyl ethanolamine, and *N*-linoleoyl ethanolamine (LEA) accompanied by increases in *N*-palmitoyl glycine (P-Gly), *N*-docosahexaenoyl glycine (D-Gly), 2-AG, and prostaglandin F2_α_ (PGF_2α_) compared to control males. Endocannabinoid levels in Crus I were significantly impacted by stress in females only, with stressed females exhibiting decreased D-Gly and 2-AG compared to normally-reared females. Accordingly, stressed females had significantly increased P-Gly and significantly decreased 2-linoleoyl glycerol (2-LG) and linoleic acid.

Striking sex and treatment differences were observed in cerebellar IP endocannabinoid levels (Figure 2, Supplementary Table 2). At baseline, females exhibited decreases in LEA, *N*-arachidonoyl taurine (A-Taur), linoleic acid, arachidonic acid (AA), and prostaglandin E_2_ (PGE_2_) compared to control males. Contrary to Crus I, treatment effects on endocannabinoids were only present in males. Stress significantly decreased 2-AG and AA in males compared to control males. In females, stress significantly increased PGE_2_. Consistent with these differences, stressed females had significantly decreased LEA and A-Taur and increased *N*-oleoyl glycine, 2-AG, and PGE_2_.

In dorsal hippocampus (Figure 3, Supplementary Table 3), control females showed significantly decreased 2-oleoyl glycerol, 2-LG, and 2-AG and increased PGF_2α_ levels compared to control males. Stressed females had significantly elevated 2-palmitoyl glycerol and 2-LG compared to stressed males. Treatment effects were only present in males, with stressed males exhibiting significantly increased PGE_2_ and PGF_2α_ compared to normally reared males.

### mRNA Analysis

Transcriptomics for this analysis are deposited in NCBI’s Gene Expression Omnibus ^59^ and are accessible through GEO Series accession number GSE139953 (https://www.ncbi.nlm.nih.gov/geo/query/acc.cgi?acc=GSE139953).

Of the 11 865 genome-wide genes evaluated, 3 009 (25.4%) had significantly different (FDR-corrected *p*<0.05) mRNA expression between all males and females; specifically, 2 057 (17.3%) significantly differed between normally reared males and females and 258 (2.2%) differed between limited bedding males and females, with 167 (1.4%) overlapping (see Fig. 4). Minimal treatment differences were observed, with 2 (0.02%) genes differing between all limited bedding and normal rearing animals; specifically, 5 (0.04%) significantly differed between normally reared and limited bedding males and 4 (0.03%) differed between normally reared and limited bedding females, with no overlap. All genes significantly differing between normally reared and limited bedding males overlapped with genes significantly differing between normally reared males and females. Two genes significantly differing between normally reared and limited bedding females overlapped with genes significantly differing between limited bedding males and females.

No significant treatment effects were observed for endocannabinoid- or cerebellar-related genes of interest within and between sex. However, five genes of interest as cannabinoid- or cerebellum-enriched genes (Fig. 4) were significantly different between normally reared males and females: females exhibited higher mRNA levels for CB1R (*Cnr1*, 1.6-fold) and interleukin 6 receptor (*Il6r*, 1.8-fold). Males exhibited higher mRNA levels for interleukin 16 (*Il16*, 1.8-fold), calcium binding protein 7 (*Cabp7*, 4.8-fold), and Purkinje cell protein 4-like 1 (*Pcp41l*, 9.7-fold). *Pcp41l* was also significantly higher (3.0-fold) in limited bedding males compared to limited bedding females.

### Behavioral Assessment

Three blinded raters scored all behaviors, except rotarod performance, which was scored digitally via the apparatus. Mean scores from all three raters were computed and used in behavioral analyses. Inter-rater reliability was deemed acceptable, with an interclass correlation of >0.95 for each behavior scored.

#### Novel Object Recognition (Fig. 5C)

A significant main effect of treatment group on discrimination ratio was observed, with limited bedding rats exhibiting decreased recognition memory (F(1,68)=5.15, *p*=0.026, ηp^2^=0.070), though performance still indicated preference for the novel object. There was no main effect of sex and no sex-treatment interaction effect. There were no significant differences by sex or treatment group in exploration time (total time interacting with the objects).

#### Social Preference (Fig. 5D)

A trending main effect of treatment was observed (F(1,40)=3.04, *p*=0.089, ηp^2^=0.071), likely due to significantly decreased social preference in females that underwent early-life stress compared to normally reared females (t(20)=2.52, *p*=0.020, d=1.074). There was no main effect of sex or sex-treatment interaction effect. There were no significant differences by sex or treatment group in exploration time (total time interacting with the object and/or social partner).

#### Rotarod (Fig. 5E)

A significant main effect in latency to fall was observed between males and females (F(1,26)=5.77, *p*=0.024, ηp^2^=0.182); specifically, males had a shorter latency to fall, likely attributed to weight differences making it more difficult for males to maintain balance on the dowels. No significant main effect was observed across treatment groups, as expected, and no sex by treatment interaction was present.

#### Elevated Plus Maze (Fig. 5F)

No main effects of sex or treatment group and no interaction effects were observed in time spent in the center, open, or closed arms. No differences were seen in open or closed arm entries (data not shown).

## Discussion

The current study is the first to our knowledge to investigate the impact of early-life stress on long-term cerebellar endocannabinoid tone in male and female rats. Notable are three major findings: (1) clear baseline sex-specific differences were observed in endocannabinoids and related lipids in a region-specific manner as well as in mRNA expression; (2) developmental stress caused sustained sex-specific changes to CNS endocannabinoid lipid tone with minimal long-term effects on mRNA; and (3) such stress impaired performance on novel object recognition and social preference.

First, baseline sex and region differences in endocannabinoid tone and mRNA expression were found, consistent with previous studies ^30,60^. Though some studies have found sex differences in cortical mRNA expression ^29^ or cerebellar regional protein differences in males only ^22^, this study, for the first time to our knowledge, identified extensive baseline sex differences in cerebellum. 2-AG was decreased in males compared to females in cerebellar IP (Fig. 3) but increased in cerebellar cortex (Crus I; Fig. 2), whereas LEA was decreased in both regions in males. Within the IP specifically, mRNA expression was significantly different between males and females at baseline (no treatment groups) across a variety of gene systems, emphasizing the robust and broad effects of sex on neural systems.

Such baseline sex differences may cause exogenous cannabinoids to differentially affect the brain in males and females. This serves as a potential explanation for the key differences in outcomes of exogenous cannabinoid exposure in animal ^61,62^ and human ^63-65^ models and perhaps also reflects sex differences in use rates ^25,64^ and adverse side effects ^66,67^ in humans. Similarly, understanding regional differences in cannabinoids may help predict directional impacts on behaviors emerging from region-specific neural circuits. Thus, baseline differences in endocannabinoid tone, as well as previous work on receptor differences ^25^, may together help make sense of differences in function, at the circuit and behavioral level.

Second, 2-AG was decreased in the cerebellar IP nucleus, but only in limited bedding males compared to normally reared males. A stress-induced rise in 2-AG during a critical developmental period may explain such a finding, resulting in a long-lasting downregulation of the endocannabinoid system to accommodate this new neuronal environment. In addition, a decrease in AA and a trending decrease in AEA was noted, suggesting that an upstream mechanism may be implicated in these effects. For example, down-regulation could be the function of a substrate imbalance, such as a decrease in phospholipids resulting in decreased overall endocannabinoid production. Although no mRNA changes for these enzymes were found, expressed protein or enzyme activity were not evaluated and, thus, this hypothesis cannot be ruled out.

No endocannabinoid changes were observed in stressed compared to non-stressed females, though stressed females did exhibit a significant increase in prostaglandin E_2_ (PGE_2_). PGE_2_ has been shown to increase cerebellar Purkinje cell arborization via estradiol production ^68-70^. Estradiol- and PGE_2_-induced cerebellar Purkinje cell arborization may function as a neuroprotective mechanism in females, perhaps further explaining the limited behavioral impairments observed in females during stress paradigms ^71-73^. Further, the increase in PGE_2_ suggests activation of cyclooxygenase (COX) enzymes, which may be related to an increase in pro-inflammatory cytokines, identified as a neural response to stress to activate microglia ^70,74^. Similarly, the increase in *N-*acyl glycines in stressed females suggests a pro-inflammatory response, as *N-*acyl glycines have been implicated in microglial migration ^74-76^. In fact, microglia have been identified as having a critical role in the sexual differentiation of the brain ^77^.

Early-life stress induced the opposite sex-specific profile in the cerebellar cortex (Fig. 2) compared to the cerebellar interpositus nucleus (Fig. 3): 2-AG was downregulated in stressed *females* compared to control females in Crus I of cerebellar cortex, whereas in cerebellar IP, 2-AG was downregulated in stressed *males* compared to control males. Cerebellar behaviors, such as associative learning, require carefully coordinated communication between the cerebellum’s nuclei and cortex. Sex-specific differences in circuit coupling resulting from endocannabinoid regulation of neuronal activation may contribute to hypotheses of baseline and stress-effects on behavioral performance differences in cerebellar tasks, like eyeblink conditioning ^47,78,79^.

Many studies looking at stress-related endocannabinoid changes have investigated the early effects of stress, finding increased 2-AG and decreased AEA ^20,80,81^. Thus, the current findings do not preclude the possibility that an acute change in endocannabinoid tone occurred immediately after the stressor; rather, the current study may suggest that any changes taking place close to the occurrence of the stressor were not enduring in these regions. Such changes may not have been maintained due to the early development of the rat hippocampus and lack of continual stressors throughout the study ^82^.

Like 2-AG, PGE_2_ showed a region-by-sex effect: upregulated in female cerebellar interpositus nucleus following stress but upregulated in stressed males in the hippocampus. As detailed above, such findings may relate to a potential pro-inflammatory response in male hippocampus following stress. Alternatively, these findings reinforce theories of foundational effects of stress on development and sexual differentiation the brain ^83^ particularly for males, further reflected by baseline regional and sex differences in the endocannabinoid system and transcriptomics generally.

By adulthood (P70), male rats having undergone stress exhibited an increase in basal systemic CORT, suggesting the stressor carries long-term effects. These findings are in line with a growing literature that males are more sensitive to stress throughout development due to sex-differences in uterine environment ^83^. In females, increased CORT was not observed, perhaps due to a resilience or compensatory mechanism contributing to female pups overcoming insults to the broader HPA system, but this resilience is lacking in the still-developing endocannabinoid system. In humans, changes in cortisol have been linked to differences in cannabis use ^84^; the increased plasticity of the HPA system may explain sex-differences in exogenous cannabinoid use and effects.

Taken together, within a given neural region, stressed males exhibited a similar lipid profile to non-stressed females. Such relationships have been previously shown in the literature and lend support to changes in estrogen receptors, perhaps explaining this feminization of neural circuits susceptible to stress in the male brain ^85^.

A third major finding is that early-life stress was associated with functional outcomes, particularly NOR performance, irrespective of sex. As shown in other studies of limited bedding ^9^ or maternal separation ^86,87^ and in the current study, rats that underwent limited bedding stress exhibited recognition memory deficits compared to controls. This was not the case for social preference, which was only impaired in stressed females in the current study. Other studies of early-life stress have shown that males are affected in social behavior later in life ^88-92^. One reason for this discrepancy may be that Crus I stress-effects on lipids were specific to females. Interestingly, a recent study by Stoodley and colleagues ^54^ found that right cerebellar Crus I disruption was related to social impairments in mice. Thus, cerebellar signaling disruption in females may have contributed to this behavioral phenotype.

### Limitations and Future Directions

Some key limitations should be addressed. First, the current findings are specific to bioactive lipid concentrations and have not yet been contextualized regarding changes in endocannabinoid-related protein expression or activity (i.e., receptor or enzyme, though major differences at the transcriptional level were not observed), though the transcriptomic data presented here will provide key insights into these pathways. Thus, the mechanistic pathway by which these endocannabinoid and prostaglandin changes have occurred is unclear and is a promising path for future inquiry. Second, although no changes in gross cerebellar integrity were observed via the rotarod task, the impacts these cerebellar changes may have on more refined cerebellar-mediated behaviors, such as eyeblink conditioning, remain unclear. The well-defined neural circuit involved in eyeblink conditioning has been shown to be impaired by other early-life stress paradigms ^47,49^, involves endocannabinoids at many steps ^93^ and is highly translatable to humans ^94^, making it an optimal task to interrogate cerebellar- and cannabinoid-specific impacts of stress ^24,95^. Third, neither estrous phase nor sex hormones were measured in this study. Excluding such measures may obscure the specific neurobiological mechanisms by which endocannabinoid and behavioral sex-differences arose, though it has been shown that cerebellar endocannabinoids do not change across the estrus cycle ^30^. Finally, it cannot be assumed that these lipid changes are solely due to the stressor; in fact, it may be that the stressor caused neuronal remodeling that altered rodent behavior, leading subsequent environmental, physiological, and social factors to impact adult lipid signaling. Thus, continued work on proximal lipid effects of stress are key.

## Conclusions

The current study provides a necessary springboard to understand sex-specific, developmental vulnerability mechanisms conferred by stress, specifically those that provide long-lasting neural changes. With increased interest in the cerebellum’s role as a node of cognitive processing ^10,96^ as well as its role in psychopathology, such as autism ^97,98^ and schizophrenia ^99^, this line of inquiry is timely and impactful. Understanding these sustained vulnerabilities may help reveal mechanisms of risk, including two-hit models of the effects of early stress and adolescent or adult cannabis use ^21,62^, and may also contribute to better-informed animal models of exogenous cannabinoid effects.

## Acknowledgements

RNA seq studies were carried out in the Center for Medical Genomics at Indiana University School of Medicine, which is partially supported by the Indiana Genomic Initiative at Indiana University (INGEN); INGEN is supported in part by the Lilly Endowment, Inc. We wish to acknowledge Dr. Andrea Hohmann and her lab for equipment use and training, including the rotarod and elevated plus maze. Additionally, we thank Jim Wager-Miller of the Mackie Lab for his training and assistance in study procedures. We also thank the IU Bloomington animal veterinary and care staff. Finally, we thank the engineering staff of the IU Psychological and Brain Sciences department, especially Jesse Goode, Rick Moore, Cameron Walker, and Allen Cody, for assistance building the apparatus for this study as well as helping set up recording equipment for behavioral tasks.

## References

1 Bradley, R. H. & Corwyn, R. F. Socioeconomic status and child development. Annual review of psychology 53, 371–399 (2002).

2 Palacios-Barrios, E. E. & Hanson, J. L. Poverty and self-regulation: Connecting psychosocial processes, neurobiology, and the risk for psychopathology. Comprehensive Psychiatry, doi:10.1016/j.comppsych.2018.12.012 (2018).

3 Anda, R. F. et al. The enduring effects of abuse and related adverse experiences in childhood. A convergence of evidence from neurobiology and epidemiology. Eur Arch Psychiatry Clin Neurosci 256, 174–186, doi:10.1007/s00406-005-0624-4 (2006).

4 Bos, K. J., Fox, N., Zeanah, C. H. & Nelson Iii, C. A. Effects of early psychosocial deprivation on the development of memory and executive function. Front Behav Neurosci 3, 16, doi:10.3389/neuro.08.016.2009 (2009).

5 Bremner, J. D., Vermetten, E., Afzal, N. & Vythilingam, M. Deficits in verbal declarative memory function in women with childhood sexual abuse-related posttraumatic stress disorder. The Journal of nervous and mental disease 192, 643–649 (2004).

6 Golier, J. A. et al. Absence of hippocampal volume differences in survivors of the Nazi Holocaust with and without posttraumatic stress disorder. Psychiatry Res 139, 53–64, doi:10.1016/j.pscychresns.2005.02.007 (2005).

7 Korkman, M., Kemp, S. L. & Kirk, U. Effects of age on neurocognitive measures of children ages 5 to 12: a cross-sectional study on 800 children from the United States. Dev Neuropsychol 20, 331–354, doi:10.1207/S15326942DN2001_2 (2001).

8 Navalta, C. P., Polcari, A., Webster, D. M., Boghossian, A. & Teicher, M. H. Effects of childhood sexual abuse on neuropsychological and cognitive function in college women. The Journal of neuropsychiatry and clinical neurosciences 18, 45–53 (2006).

9 Brunson, K. L. et al. Mechanisms of late-onset cognitive decline after early-life stress. J Neurosci 25, 9328–9338, doi:10.1523/JNEUROSCI.2281-05.2005 (2005).

10 Buckner, R. L. The cerebellum and cognitive function: 25 years of insight from anatomy and neuroimaging. Neuron 80, 807–815, doi:10.1016/j.neuron.2013.10.044 (2013).

11 Sokolov, A. A. The Cerebellum in Social Cognition. Frontiers in Cellular Neuroscience 12, doi:10.3389/fncel.2018.00145 (2018).

12 Sokolov, A. A., Miall, R. C. & Ivry, R. B. The Cerebellum: Adaptive Prediction for Movement and Cognition. Trends Cogn Sci 21, 313–332, doi:10.1016/j.tics.2017.02.005 (2017).

13 Stoodley, C. J. & Schmahmann, J. D. Evidence for topographic organization in the cerebellum of motor control versus cognitive and affective processing. Cortex 46, 831–844, doi:10.1016/j.cortex.2009.11.008 (2010).

14 Stoodley, C. J., Valera, E. M. & Schmahmann, J. D. Functional topography of the cerebellum for motor and cognitive tasks: an fMRI study. Neuroimage 59, 1560–1570, doi:10.1016/j.neuroimage.2011.08.065 (2012).

15 Charil, A., Laplante, D. P., Vaillancourt, C. & King, S. Prenatal stress and brain development. Brain Res Rev 65, 56–79, doi:10.1016/j.brainresrev.2010.06.002 (2010).

16 Popoli, M., Yan, Z., McEwen, B. S. & Sanacora, G. The stressed synapse: the impact of stress and glucocorticoids on glutamate transmission. Nat Rev Neurosci 13, 22–37, doi:10.1038/nrn3138 (2011).

17 McEwen, M., Johnson, P., Neatherlin, J., Millard, M. W. & Lawrence, G. School-based management of chronic asthma among inner-city African-American schoolchildren in Dallas, Texas. J Sch Health 68, 196–201 (1998).

18 Glaser, D. Child abuse and neglect and the brain--a review. J Child Psychol Psychiatry 41, 97–116 (2000).

19 Hill, M. N. & Tasker, J. G. Endocannabinoid signaling, glucocorticoid-mediated negative feedback, and regulation of the hypothalamic-pituitary-adrenal axis. Neuroscience 204, 5–16, doi:10.1016/j.neuroscience.2011.12.030 (2012).

20 Morena, M., Patel, S., Bains, J. S. & Hill, M. N. Neurobiological Interactions Between Stress and the Endocannabinoid System. Neuropsychopharmacology 41, 80–102, doi:10.1038/npp.2015.166 (2016).

21 Saravia, R. et al. Concomitant THC and stress adolescent exposure induces impaired fear extinction and related neurobiological changes in adulthood. Neuropharmacology 144, 345–357, doi:10.1016/j.neuropharm.2018.11.016 (2019).

22 Suarez, J. et al. Immunohistochemical description of the endogenous cannabinoid system in the rat cerebellum and functionally related nuclei. J Comp Neurol 509, 400–421, doi:10.1002/cne.21774 (2008).

23 Mackie, K. Cannabinoid receptors: where they are and what they do. J Neuroendocrinol 20 Suppl 1, 10–14, doi:10.1111/j.1365-2826.2008.01671.x (2008).

24 Steinmetz, A. B. & Freeman, J. H. Central cannabinoid receptors modulate acquisition of eyeblink conditioning. Learn Mem 17, 571–576, doi:10.1101/lm.1954710 (2010).

25 Viveros, M. P., Marco, E. M., Lopez-Gallardo, M., Garcia-Segura, L. M. & Wagner, E. J. Framework for sex differences in adolescent neurobiology: a focus on cannabinoids. Neurosci Biobehav Rev 35, 1740–1751, doi:10.1016/j.neubiorev.2010.09.005 (2011).

26 Goel, N., Workman, J. L., Lee, T. T., Innala, L. & Viau, V. Sex differences in the HPA axis. Compr Physiol 4, 1121–1155, doi:10.1002/cphy.c130054 (2014).

27 Vangopoulou, C. et al. Effects of an early life experience on rat brain cannabinoid receptors in adolescence and adulthood. IBRO Reports 5, 1–9, doi:10.1016/j.ibror.2018.05.002 (2018).

28 Dow-Edwards, D., Frank, A., Wade, D., Weedon, J. & Izenwasser, S. Sexually-dimorphic alterations in cannabinoid receptor density depend upon prenatal/early postnatal history. Neurotoxicol Teratol 58, 31–39, doi:10.1016/j.ntt.2016.09.004 (2016).

29 Marco, E. M. et al. Consequences of early life stress on the expression of endocannabinoid-related genes in the rat brain. Behav Pharmacol 25, 547–556, doi:10.1097/FBP.0000000000000068 (2014).

30 Bradshaw, H. B., Rimmerman, N., Krey, J. F. & Walker, J. M. Sex and hormonal cycle differences in rat brain levels of pain-related cannabimimetic lipid mediators. Am J Physiol Regul Integr Comp Physiol 291, R349–358, doi:10.1152/ajpregu.00933.2005 (2006).

31 Herman, J. P. & Cullinan, W. E. Neurocircuitry of stress: central control of the hypothalamo–pituitary–adrenocortical axis. Trends in neurosciences 20, 78–84 (1997).

32 Hutcheson, N. L. et al. Effective connectivity during episodic memory retrieval in schizophrenia participants before and after antipsychotic medication. Hum Brain Mapp 36, 1442–1457, doi:10.1002/hbm.22714 (2015).

33 Herman, J. P. Regulation of adrenocorticosteroid receptor mRNA expression in the central nervous system. Cellular and molecular neurobiology 13, 349–372 (1993).

34 McEwen, B. S. in Adrenal actions on brain 1–22 (Springer, 1982).

35 Bremner, J. D. et al. Magnetic resonance imaging-based measurement of hippocampal volume in posttraumatic stress disorder related to childhood physical and sexual abuse— a preliminary report. Biological psychiatry 41, 23–32 (1997).

36 Bremner, J. D. et al. MRI and PET study of deficits in hippocampal structure and function in women with childhood sexual abuse and posttraumatic stress disorder. American Journal of Psychiatry 160, 924–932 (2003).

37 Avishai□Eliner, S., Gilles, E. E., Eghbal□Ahmadi, M., Bar□El, Y. & Baram, T. Z. Altered regulation of gene and protein expression of hypothalamic□pituitary□adrenal axis components in an immature rat model of chronic stress. Journal of neuroendocrinology 13, 799–807 (2001).

38 Youssef, M. et al. Early life stress delays hippocampal development and diminishes the adult stem cell pool in mice. Sci Rep 9, 4120, doi:10.1038/s41598-019-40868-0 (2019).

39 Zorn, J. V. et al. Cortisol stress reactivity across psychiatric disorders: a systematic review and meta-analysis. Psychoneuroendocrinology 77, 25–36 (2017).

40 Altman, J. Morphological development of the rat cerebellum and some of its mechanisms. Experimental Brain Research Supplemental 6, 8–49 (1982).

41 Arpino, C. et al. Preterm birth and neurodevelopmental outcome: a review. Childs Nerv Syst 26, 1139–1149, doi:10.1007/s00381-010-1125-y (2010).

42 Limperopoulos, C. Late Gestation Cerebellar Growth Is Rapid and Impeded by Premature Birth. Pediatrics 115, 688–695, doi:10.1542/peds.2004-1169 (2005).

43 Brooks, S. J. et al. Childhood adversity is linked to differential brain volumes in adolescents with alcohol use disorder: a voxel-based morphometry study. Metab Brain Dis 29, 311–321, doi:10.1007/s11011-014-9489-4 (2014).

44 Brooks, S. J. et al. Early-life adversity and orbitofrontal and cerebellar volumes in adults with obsessive-compulsive disorder: voxel-based morphometry study. Br J Psychiatry 208, 34–41, doi:10.1192/bjp.bp.114.162610 (2016).

45 De Bellis, M. D. & Kuchibhatla, M. Cerebellar volumes in pediatric maltreatment-related posttraumatic stress disorder. Biol Psychiatry 60, 697–703, doi:10.1016/j.biopsych.2006.04.035 (2006).

46 Llorente, R. et al. Early maternal deprivation in rats induces gender-dependent effects on developing hippocampal and cerebellar cells. Int J Dev Neurosci 27, 233–241, doi:10.1016/j.ijdevneu.2009.01.002 (2009).

47 Wilber, A. A., Southwood, C. J., Sokoloff, G., Steinmetz, J. E. & Wellman, C. L. Neonatal maternal separation alters adult eyeblink conditioning and glucocorticoid receptor expression in the interpositus nucleus of the cerebellum. Dev Neurobiol 67, 1751–1764, doi:10.1002/dneu.20549 (2007).

48 Wilber, A. A. & Wellman, C. L. Neonatal maternal separation alters the development of glucocorticoid receptor expression in the interpositus nucleus of the cerebellum. Int J Dev Neurosci 27, 649–654, doi:10.1016/j.ijdevneu.2009.08.001 (2009).

49 Wilber, A. A. & Wellman, C. L. Neonatal maternal separation-induced changes in glucocorticoid receptor expression in posterior interpositus interneurons but not projection neurons predict deficits in adult eyeblink conditioning. Neurosci Lett 460, 214–218, doi:10.1016/j.neulet.2009.05.076 (2009).

50 Parker, K. L. et al. Delta-frequency stimulation of cerebellar projections can compensate for schizophrenia-related medial frontal dysfunction. Mol Psychiatry 22, 647–655, doi:10.1038/mp.2017.50 (2017).

51 McAfee, S. S., Liu, Y., Sillitoe, R. V. & Heck, D. H. Cerebellar Lobulus Simplex and Crus I Differentially Represent Phase and Phase Difference of Prefrontal Cortical and Hippocampal Oscillations. Cell Rep 27, 2328–2334 e2323, doi:10.1016/j.celrep.2019.04.085 (2019).

52 Shipman, M. L. & Green, J. T. Cerebellum and cognition: Does the rodent cerebellum participate in cognitive functions? Neurobiol Learn Mem, doi:10.1016/j.nlm.2019.02.006 (2019).

53 Deverett, B., Koay, S. A., Oostland, M. & Wang, S. S. Cerebellar involvement in an evidence-accumulation decision-making task. Elife 7, doi:10.7554/eLife.36781 (2018).

54 Stoodley, C. J. et al. Altered cerebellar connectivity in autism and cerebellar-mediated rescue of autism-related behaviors in mice. Nat Neurosci 20, 1744–1751, doi:10.1038/s41593-017-0004-1 (2017).

55 Baker, L. M. et al. Impact of early vs. late childhood early life stress on brain morphometrics. Brain Imaging Behav 7, 196–203, doi:10.1007/s11682-012-9215-y (2013).

56 Bonne, O. et al. Longitudinal MRI study of hippocampal volume in trauma survivors with PTSD. American Journal of Psychiatry 158, 1248–1251 (2001).

57 Molet, J., Maras, P. M., Avishai-Eliner, S. & Baram, T. Z. Naturalistic rodent models of chronic early-life stress. Dev Psychobiol 56, 1675–1688, doi:10.1002/dev.21230 (2014).

58 Leishman, E., Mackie, K., Luquet, S. & Bradshaw, H. B. Lipidomics profile of a NAPE-PLD KO mouse provides evidence of a broader role of this enzyme in lipid metabolism in the brain. Biochim Biophys Acta 1861, 491–500, doi:10.1016/j.bbalip.2016.03.003 (2016).

59 Edgar, R., Domrachev, M. & Lash, A. E. Gene Expression Omnibus: NCBI gene expression and hybridization array data repository. Nucleic acids research 30, 207–210 (2002).

60 Wagner, E. J. Sex differences in cannabinoid-regulated biology: A focus on energy homeostasis. Front Neuroendocrinol 40, 101–109, doi:10.1016/j.yfrne.2016.01.003 (2016).

61 Becker, J. B. & Koob, G. F. Sex Differences in Animal Models: Focus on Addiction. Pharmacol Rev 68, 242–263, doi:10.1124/pr.115.011163 (2016).

62 Viveros, M. P. et al. The endocannabinoid system in critical neurodevelopmental periods: sex differences and neuropsychiatric implications. J Psychopharmacol 26, 164–176, doi:10.1177/0269881111408956 (2012).

63 Crane, N. A., Schuster, R. M., Fusar-Poli, P. & Gonzalez, R. Effects of cannabis on neurocognitive functioning: recent advances, neurodevelopmental influences, and sex differences. Neuropsychol Rev 23, 117–137, doi:10.1007/s11065-012-9222-1 (2013).

64 Ketcherside, A., Baine, J. & Filbey, F. Sex Effects of Marijuana on Brain Structure and Function. Curr Addict Rep 3, 323–331, doi:10.1007/s40429-016-0114-y (2016).

65 Rubino, T. & Parolaro, D. Sexually dimorphic effects of cannabinoid compounds on emotion and cognition. Front Behav Neurosci 5, 64, doi:10.3389/fnbeh.2011.00064 (2011).

66 Bonnet, U. & Preuss, U. W. The cannabis withdrawal syndrome: current insights. Subst Abuse Rehabil 8, 9–37, doi:10.2147/SAR.S109576 (2017).

67 Fattore, L. & Fratta, W. How important are sex differences in cannabinoid action? Br J Pharmacol 160, 544–548, doi:10.1111/j.1476-5381.2010.00776.x (2010).

68 Dean, S. L. & McCarthy, M. M. Steroids, sex and the cerebellar cortex: implications for human disease. Cerebellum 7, 38–47, doi:10.1007/s12311-008-0003-6 (2008).

69 Dean, S. L. et al. Prostaglandin E2 stimulates estradiol synthesis in the cerebellum postnatally with associated effects on Purkinje neuron dendritic arbor and electrophysiological properties. Endocrinology 153, 5415–5427, doi:10.1210/en.2012-1350 (2012).

70 Hoffman, J. F., Wright, C. L. & McCarthy, M. M. A Critical Period in Purkinje Cell Development Is Mediated by Local Estradiol Synthesis, Disrupted by Inflammation, and Has Enduring Consequences Only for Males. J Neurosci 36, 10039–10049, doi:10.1523/JNEUROSCI.1262-16.2016 (2016).

71 Bale, T. L. et al. Early life programming and neurodevelopmental disorders. Biol Psychiatry 68, 314–319, doi:10.1016/j.biopsych.2010.05.028 (2010).

72 Leuner, B., Mendolia-Loffredo, S. & Shors, T. J. High levels of estrogen enhance associative memory formation in ovariectomized females. Psychoneuroendocrinology 29, 883–890, doi:10.1016/j.psyneuen.2003.08.001 (2004).

73 Wright, C. L., Hoffman, J. H. & McCarthy, M. M. Evidence that inflammation promotes estradiol synthesis in human cerebellum during early childhood. Transl Psychiatry 9, 58, doi:10.1038/s41398-018-0363-8 (2019).

74 Lenz, K. M. & McCarthy, M. M. A starring role for microglia in brain sex differences. Neuroscientist 21, 306–321, doi:10.1177/1073858414536468 (2015).

75 Bradshaw, H. B. et al. The endocannabinoid anandamide is a precursor for the signaling lipid N-arachidonoyl glycine by two distinct pathways. BMC Biochem 10, 14, doi:10.1186/1471-2091-10-14 (2009).

76 McHugh, D., Roskowski, D., Xie, S. & Bradshaw, H. B. Delta(9)-THC and N-arachidonoyl glycine regulate BV-2 microglial morphology and cytokine release plasticity: implications for signaling at GPR18. Front Pharmacol 4, 162, doi:10.3389/fphar.2013.00162 (2014).

77 VanRyzin, J. W., Pickett, L. A. & McCarthy, M. M. Microglia: Driving critical periods and sexual differentiation of the brain. Dev Neurobiol 78, 580–592, doi:10.1002/dneu.22569 (2018).

78 Dalla, C. & Shors, T. J. Sex differences in learning processes of classical and operant conditioning. Physiol Behav 97, 229–238, doi:10.1016/j.physbeh.2009.02.035 (2009).

79 Wood, G. E. & Shors, T. J. Stress facilitates classical conditioning in males, but impairs classical conditioning in females through activational effects of ovarian hormones. Proceedings of the National Academy of Sciences 95, 4066–4071 (1998).

80 Gorzalka, B. B., Hill, M. N. & Hillard, C. J. Regulation of endocannabinoid signaling by stress: implications for stress-related affective disorders. Neurosci Biobehav Rev 32, 1152–1160, doi:10.1016/j.neubiorev.2008.03.004 (2008).

81 Patel, S., Kingsley, P. J., Mackie, K., Marnett, L. J. & Winder, D. G. Repeated homotypic stress elevates 2-arachidonoylglycerol levels and enhances short-term endocannabinoid signaling at inhibitory synapses in basolateral amygdala. Neuropsychopharmacology 34, 2699–2709, doi:10.1038/npp.2009.101 (2009).

82 Bayer, S. A. Development of the hippocampal region in the rat II. Morphogenesis during embryonic and early postnatal life. Journal of Comparative Neurology 190, 115–134 (1980).

83 McCarthy, M. M. in Oxford Research Encyclopedia of Neuroscience (2019).

84 Huizink, A. C., Ferdinand, R. F., Ormel, J. & Verhulst, F. C. Hypothalamic-pituitary-adrenal axis activity and early onset of cannabis use. Addiction 101, 1581–1588, doi:10.1111/j.1360-0443.2006.01570.x (2006).

85 Margaret McCarthy, M. M. & Nugent, B. Epigenetic influences on the developing brain: effects of hormones and nutrition. Advances in Genomics and Genetics, doi:10.2147/agg.S58625 (2015).

86 Aisa, B., Tordera, R., Lasheras, B., Del Rio, J. & Ramirez, M. J. Cognitive impairment associated to HPA axis hyperactivity after maternal separation in rats. Psychoneuroendocrinology 32, 256–266, doi:10.1016/j.psyneuen.2006.12.013 (2007).

87 Hulshof, H. J. et al. Maternal separation decreases adult hippocampal cell proliferation and impairs cognitive performance but has little effect on stress sensitivity and anxiety in adult Wistar rats. Behav Brain Res 216, 552–560, doi:10.1016/j.bbr.2010.08.038 (2011).

88 Rincon-Cortes, M. & Sullivan, R. M. Emergence of social behavior deficit, blunted corticolimbic activity and adult depression-like behavior in a rodent model of maternal maltreatment. Transl Psychiatry 6, e930, doi:10.1038/tp.2016.205 (2016).

89 Raineki, C., Moriceau, S. & Sullivan, R. M. Developing a neurobehavioral animal model of infant attachment to an abusive caregiver. Biol Psychiatry 67, 1137–1145, doi:10.1016/j.biopsych.2009.12.019 (2010).

90 Raineki, C., Cortes, M. R., Belnoue, L. & Sullivan, R. M. Effects of early-life abuse differ across development: infant social behavior deficits are followed by adolescent depressive-like behaviors mediated by the amygdala. J Neurosci 32, 7758–7765, doi:10.1523/JNEUROSCI.5843-11.2012 (2012).

91 Dalmaz, C., Noschang, C., Krolow, R., Raineki, C. & Lucion, A. B. How postnatal insults may program development: studies in animal models. Adv Neurobiol 10, 121–147, doi:10.1007/978-1-4939-1372-5_7 (2015).

92 Guadagno, A., Wong, T. P. & Walker, C. D. Morphological and functional changes in the preweaning basolateral amygdala induced by early chronic stress associate with anxiety and fear behavior in adult male, but not female rats. Prog Neuropsychopharmacol Biol Psychiatry 81, 25–37, doi:10.1016/j.pnpbp.2017.09.025 (2018).

93 Tabata, T. & Kano, M. in Handbook of Neurochemistry and Molecular Neurobiology Ch. Chapter 6, 63–86 (2009).

94 Reeb-Sutherland, B. C. & Fox, N. A. Eyeblink conditioning: a non-invasive biomarker for neurodevelopmental disorders. J Autism Dev Disord 45, 376–394, doi:10.1007/s10803-013-1905-9 (2015).

95 Steinmetz, J. E. Brain substrates of classical eyeblink conditioning: a highly localized but also distributed system. Behavioural Brain Research 110, 13–24, doi:10.1016/s0166-4328(99)00181-3 (2000).

96 Schmahmann, J. D. The cerebellum and cognition. Neurosci Lett 688, 62–75, doi:10.1016/j.neulet.2018.07.005 (2019).

97 Kern, J. K. The possible role of the cerebellum in autism/PDD: disruption of a multisensory feedback loop. Medical Hypotheses 59, 255–260, doi:10.1016/s0306-9877(02)00212-8 (2002).

98 Wang, S. S., Kloth, A. D. & Badura, A. The cerebellum, sensitive periods, and autism. Neuron 83, 518–532, doi:10.1016/j.neuron.2014.07.016 (2014).

99 Picard, H., Amado, I., Mouchet-Mages, S., Olie, J. P. & Krebs, M. O. The role of the cerebellum in schizophrenia: an update of clinical, cognitive, and functional evidences. Schizophr Bull 34, 155–172, doi:10.1093/schbul/sbm049 (2008).

