## Supplemental Tables for "Long-term aberrations to cerebellar endocannabinoids induced by early-life stress"

**Supplementary Table 1: Crus/HVI Endocannabinoids**

| Lipid Species | Male | | Female | | |
|---|---|---|---|---|---|
| | **Normal Rearing** (N=8) | **Limited Bedding** (N=8) | **Normal Rearing** (N=6) | **Limited Bedding** (N=7) | ***p*-value*** |
| ***N*-acyl ethanolamine** | | | | | |
| *N*-palmitoyl ethanolamine | 4.9E-10± 6.25E-11 | 5.52E-10± 2.86E-11 | 3.77E-10± 5.46E-11 | 4.64E-10± 5.94E-11 | NS |
| *N*-stearoyl ethanolamine | *2.41E-10± 3.97E-11* | 2.34E-10± 2.83E-11 | *1.21E-10± 2.52E-11* | 2.11E-10± 4.03E-11 | *0.028* |
| *N*-oleoyl ethanolamine | *2.1E-09± 3.28E-10* | 1.96E-09± 3.31E-10 | *1.03E-09± 2.09E-10* | 1.66E-09± 3.17E-10 | *0.027* |
| *N*-linoleoyl ethanolamine | *2.01E-11± 1.93E-12* | 1.72E-11± 1.58E-12 | *9.66E-12± 1.24E-12* | 1.45E-11± 1.78E-12 | *<0.001* |
| *N*-arachidonoyl ethanolamine | 1.79E-11± 2.91E-12 | 1.25E-11± 5.37E-13 | 1.58E-11± 2.34E-12 | 1.62E-11± 2.59E-12 | NS |
| *N*-docosahexaenoyl ethanolamine | 3.16E-11± 2.66E-12 | 3.06E-11± 9.74E-13 | 3.01E-11± 4.28E-12 | 3.04E-11± 4.26E-12 | NS |
| ***N*-acyl glycine** | | | | | |
| *N*-palmitoyl glycine | *4.85E-11± 3.99E-12* | **4.68E-11± 3.65E-12** | *6.46E-11± 7.94E-12* | **6.09E-11± 2.64E-12** | *0.021*<br>**0.045** |
| *N*-stearoyl glycine | 1..45E-11± 9.22E-13 | 1.4E-11± 9.11E-13 | 1.5E-11± 1.19E-12 | 1.64E-11± 1.42E-12 | NS |
| *N*-oleoyl glycine | 4.35E-12± 5.74E-13 | 3.6E-12± 2.47E-13 | 3.19E-12± 3.06E-13 | 4.26E-12± 4.51E-13 | NS |
| *N*-linoleoyl glycine | 6.95E-13± 8.63E-14 | 8E-13± 1.73E-13 | 8.49E-13± 2.43E-13 | 7.63E-13± 1.39E-13 | NS |
| *N*-arachidonoyl glycine | 7.09E-12± 2.22E-13 | 5.83E-12± 8.92E-14 | 6.48E-12± 8.35E-13 | 6.23E-12± 7.1E-13 | NS |
| *N*-docosahexaenoyl glycine | *1.5E-12± 1.22E-13* | 1.61E-12± 3.09E-13 | *2.08E-12± 8.27E-14♀* | 1.46E-12± 1.16E-13♀ | *0.005*<br>0.004♀ |
| ***N*-acyl taurine** | | | | | |
| *N*-palmitoyl taurine | 1.25E-10± 8.87E-12 | 1.16E-10± 8.16E-12 | 1.22E-10± 3.31E-12 | 1.26E-10± 6.23E-12 | NS |
| *N*-stearoyl taurine | 1.08E-10± 7.84E-12 | 1.26E-10± 9.89E-12 | 1.34E-10± 1.32E-11 | 1.16E-10± 6.49E-12 | NS |
| *N*-oleoyl taurine | 2.55E-11± 2.25E-12 | 2.04E-11± 2.33E-12 | 2.2E-11± 3.83E-12 | 2.17E-11± 2.48E-12 | NS |
| *N*-arachidonoyl taurine | 2.05E-10± 1.65E-11 | 2.22E-10± 1.6E-11 | 2.06E-10± 1.61E-11 | 2.05E-10± 1.15E-11 | NS |
| **2-acyl glycerol** | | | | | |
| 2-palmitoyl glycerol | 3.37E-09± 5.2E-10 | 4.41E-09± 1.04E-09 | 4.69E-09± 1.28E-09 | 2.91E-09± 4.66E-10 | NS |
| 2-oleoyl glycerol | 6.36E-09± 6.41E-10 | 6.29E-09± 5.2E-10 | 5.89E-09± 6.02E-10 | 4.88E-09± 5.23E-10 | NS |
| 2-linoleoyl glycerol | 6.48E-10± 3.42E-11 | **7.01E-10± 6.9E-11** | 6.31E-10± 8.07E-11 | **4.96E-10± 5.14E-11** | **0.020** |
| 2-arachidonoyl glycerol | *1.22E-09± 3.6E-11* | 1.29E-09± 6.91E-11 | *1.26E-09± 5.1E-11♀* | 1.09E-09± 1.18E-10♀ | *0.020*<br>0.002♀ |
| **Free Fatty Acids** | | | | | |

| | | | | | |
|---|---|---|---|---|---|
| Oleic acid | 3.84E-10± 4.09E-11 | 4.13E-10± 3.95E-11 | 3.23E-10± 3.32E-11 | 3.33E-10± 4.55E-11 | NS |
| Linoleic acid | 6.07E-10± 4.26E-11 | **6.25E-10± 2.93E-11** | 5.11E-10± 6.17E-11 | **4.87E-10± 5.66E-11** | **0.047** |
| Arachidonic acid | 7.23E-09± 4.26E-10 | 8E-09± 4.43E-10 | 7.65E-09± 5.65E-10 | 7.48E-09± 6.51E-10 | NS |
| **Prostaglandins** | | | | | |
| $PGE_2$ | 9.54E-11± 7.62E-12 | 8.29E-11± 9.18E-12 | 8.18E-11± 6.88E-12 | 6.82E-11± 6.51E-12 | NS |
| $PGF_{2\alpha}$ | *1.66E-10± 1.34E-11* | 2.04E-10± 7.92E-12 | *2.25E-10± 1.81E-11* | 1.96E-10± 2.33E-11 | *0.017* |
| 6-keto$PGF_{1\alpha}$ | BAL | BAL | BAL | BAL | -- |
| **Sample Mass** | | | | | |
| Sample Mass | *0.02±1.12E-3* | 0.02±7.3E-4 | *0.02±2.1E-3* | 0.02±8.1E-4 | *0.001* |

Data are moles per gram tissue and are shown as means ± SE. Values in light face have no significant difference among the groups. *Least Significant Difference corrected $p \leq 0.05$. *Italicized* values denote a significant sex effect for normally reared animals, whereas those in **bold** denote a significant sex effect for limited bedding animals; ♀=significant treatment effect for females, NS=Not Significant. BAL=Below Analytical Limits.

**Supplementary Table 2: Interpositus Nucleus Endocannabinoids**

| Lipid Species | Male | | Female | | |
|---|---|---|---|---|---|
| | **Normal Rearing** (N=8) | **Limited Bedding** (N=8) | **Normal Rearing** (N=6) | **Limited Bedding** (N=7) | ***p*-value*** |
| ***N*-acyl ethanolamine** | | | | | |
| *N*-palmitoyl ethanolamine | 2.81E-10± 3.18E-11 | 3.29E-10± 3.87E-11 | 3.13E-10± 4.37E-11 | 3.46E-10± 3.11E-11 | NS |
| *N*-stearoyl ethanolamine | 7.45E-11± 1.14E-11 | 7.31E-11± 7.65E-12 | 7.17E-11± 5.79E-12 | 9.10E-11± 1.31E-11 | NS |
| *N*-oleoyl ethanolamine | 2.25E-10± 2.86E-11 | 2.24E-10± 2.03E-11 | 2.10E-10± 2.37E-11 | 2.35E-10± 2.84E-11 | NS |
| *N*-linoleoyl ethanolamine | *2.93E-11± 2.86E-12* | 2.75E-11± 1.85E-12 | *2.03E-11± 2.02E-12* | 2.10E-11± 2.71E-12 | *0.020* |
| *N*-arachidonoyl ethanolamine | 1.47E-11± 1.60E-12 | 1.11E-11± 3.84E-13 | 1.10E-11± 1.00E-12 | 1.16E-11± 2.15E-12 | NS |
| *N*-docosahexaenoyl ethanolamine | 2.57E-11± 2.87E-12 | 2.29E-11± 1.79E-12 | 2.08E-11± 2.25E-12 | 2.25E-11± 2.36E-12 | NS |
| ***N*-acyl glycine** | | | | | |
| *N*-palmitoyl glycine | 1.39E-11± 1.14E-12 | 1.29E-11± 1.53E-12 | 1.50E-11± 2.88E-12 | 1.81E-11± 1.98E-12 | NS |
| *N*-stearoyl glycine | 2.42E-12± 1.82E-13 | 2.41E-12± 4.29E-13 | 2.39E-12± 6.30E-13 | 3.24E-12± 4.56E-13 | NS |
| *N*-oleoyl glycine | 2.94E-12± 2.31E-13 | **2.47E-12± 1.74E-13** | 3.56E-12± 5.58E-13 | **3.65E-12± 5.59E-13** | **0.038** |
| *N*-linoleoyl glycine | 9.89E-13± 1.35E-13 | 8.34E-13± 1.65E-13 | 1.26E-12± 3.23E-13 | 1.01E-12± 1.12E-13 | NS |
| *N*-arachidonoyl glycine | 6.71E-12± 9.91E-13 | 5.69E-12± 4.78E-13 | 5.37E-12± 8.62E-13 | 6.62E-12± 1.47E-12 | NS |
| *N*-docosahexaenoyl glycine | 2.55E-12± 2.40E-13 | 2.01E-12± 1.92E-13 | 2.78E-12± 2.76E-13 | 3.47E-12± 5.52E-13 | NS |
| ***N*-acyl taurine** | | | | | |
| *N*-palmitoyl taurine | 1.57E-10± 9.45E-12 | 1.51E-10± 7.65E-12 | 1.57E-10± 1.56E-11 | 1.44E-10± 7.70E-12 | NS |
| *N*-stearoyl taurine | 1.72E-10± 4.10E-12 | 1.82E-10± 1.39E-11 | 1.78E-10± 1.24E-11 | 1.89E-10± 1.26E-11 | NS |
| *N*-oleoyl taurine | 2.79E-11± 2.86E-12 | 3.49E-11± 5.21E-12 | 3.04E-11± 3.22E-12 | 3.59E-11± 4.54E-12 | NS |
| *N*-arachidonoyl taurine | *2.53E-11± 7.46E-12* | **2.53E-11± 7.42E-12** | *1.68E-11± 3.42E-12* | **1.59E-11± 5.18E-12** | *0.017* **0.009** |
| **2-acyl glycerol** | | | | | |
| 2-palmitoyl glycerol | 2.85E-08± 8.60E-09 | 2.92E-08± 1.11E-08 | 2.56E-08± 1.21E-08 | 4.05E-08± 1.67E-08 | NS |
| 2-oleoyl glycerol | 9.81E-09± 1.44E-09 | 7.79E-09± 9.90E-10 | 8.85E-09± 1.24E-09 | 9.65E-09± 1.62E-09 | NS |
| 2-linoleoyl glycerol | 9.45E-10± 1.14E-10 | 7.13E-10± 5.86E-11 | 7.95E-10± 7.76E-11 | 9.00E-10± 7.54E-11 | NS |
| 2-arachidonoyl glycerol | 1.49E-09± 1.65E-10♂ | **1.07E-09± 7.53E-11♂** | 1.54E-09± 9.19E-11 | **1.80E-09± 8.97E-11** | 0.015♂ **<0.001** |
| **Free Fatty Acids** | | | | | |

| | | | | | |
|---|---|---|---|---|---|
| Oleic acid | 2.69E-10±<br>2.41E-11 | 2.56E-10±<br>2.69E-11 | 2.24E-10±<br>1.51E-11 | 2.62E-10±<br>3.89E-11 | NS |
| Linoleic acid | *6.04E-13±<br>7.17E-14* | 4.44E-13±<br>6.41E-14 | *3.35E-13±<br>6.12E-14* | 2.70E-13±<br>4.90E-14 | *0.008* |
| Arachidonic acid | *4.35E-10±<br>2.05E-11*♂ | 3.58E-10±<br>2.28E-11♂ | *3.72E-10±<br>1.12E-11* | 3.34E-10±<br>2.38E-11 | *0.049*<br>0.011♂ |
| **Prostaglandins** | | | | | |
| $PGE_2$ | *8.93E-11±<br>6.77E-12* | **7.57E-11±<br>4.02E-12** | *6.56E-11±<br>7.07E-12*♀ | **9.93E-11±<br>9.35E-12**♀ | *0.026*<br>**0.024**<br>0.004♀ |
| $PGF_{2\alpha}$ | BAL | BAL | BAL | BAL | |
| 6-keto$PGF_{1\alpha}$ | 3.18E-11±<br>4.66E-12 | 3.72E-11±<br>4.04E-12 | 4.07E-11±<br>6.82E-12 | 4.79E-11±<br>7.83E-12 | NS |
| **Sample Mass** | | | | | |
| Sample Mass | *0.02±5.03E-4* | **0.02±6.34E-4** | *0.02±9.30E-4* | **0.02±1.02E-3** | *0.020*<br>**0.001** |

Data are moles per gram tissue and are shown as means ± SE. Values in light face have no significant difference among the groups. *Least Significant Difference corrected $p \leq 0.05$. *Italicized* values denote a significant sex effect for normally reared animals, whereas those in **bold** denote a significant sex effect for limited bedding animals; ♂=significant treatment effect for males, ♀=significant treatment effect for females, NS=Not Significant. BAL=Below Analytical Limits.

**Supplementary Table 3: Dorsal Hippocampus Endocannabinoids**

| Lipid Species | Male | | Female | | |
|---|---|---|---|---|---|
| | **Normal Rearing** (N=8) | **Limited Bedding** (N=8) | **Normal Rearing** (N=6) | **Limited Bedding** (N=7) | ***p*-value*** |
| ***N*-acyl ethanolamine** | | | | | |
| *N*-palmitoyl ethanolamine | 5.62E-10± 7.69E-11 | 6.47E-10± 1.16E-10 | 3.55E-10± 4.5E-11 | 4.12E-10± 8.05E-11 | NS |
| *N*-stearoyl ethanolamine | 5.2E-11± 1.07E-11 | 7.91E-11± 9.59E-12 | 4.53E-11± 8.46E-12 | 5.16E-11± 1.16E-11 | NS |
| *N*-oleoyl ethanolamine | 6.36E-10± 4.76E-11 | 7.61E-10± 7.31E-11 | 4.73E-10± 3.82E-11 | 6.39E-10± 1.3E-10 | NS |
| *N*-linoleoyl ethanolamine | 1.07E-10± 1.29E-11 | 1.04E-10± 1.28E-11 | 8.61E-11± 1.26E-11 | 9.44E-11± 1.28E-11 | NS |
| *N*-arachidonoyl ethanolamine | 4.66E-11± 7.07E-12 | 4.21E-11± 5.11E-12 | 4.15E-11± 8.57E-12 | 4.81E-11± 7.18E-12 | NS |
| *N*-docosahexaenoyl ethanolamine | 3.83E-11± 4.87E-12 | 4.01E-11± 5.47E-12 | 3.1E-11± 5.29E-12 | 4.47E-11± 8.96E-12 | NS |
| ***N*-acyl glycine** | | | | | |
| *N*-palmitoyl glycine | 1.15E-11± 6.07E-13 | 1.5E-11± 9.45E-13 | 1.4E-11± 2.27E-12 | 1.29E-11± 1.57E-12 | NS |
| *N*-stearoyl glycine | 6.75E-12± 4.34E-13 | 6.63E-12± 4.92E-13 | 6.85E-12± 5E-13 | 6.97E-12± 3.85E-13 | NS |
| *N*-oleoyl glycine | 5.28E-12± 3.47E-13 | 6.2E-12± 2.66E-13 | 5.57E-12± 3.36E-13 | 6.25E-12± 3.89E-13 | NS |
| *N*-linoleoyl glycine | 4.72E-13± 4.11E-14 | 5.97E-13± 5.08E-14 | 6.15E-13± 8.04E-14 | 5.81E-13± 9.82E-14 | NS |
| *N*-arachidonoyl glycine | 1.15E-11± 1E-12 | 1.15E-11± 8.6E-13 | 1.23E-11± 9.05E-13 | 1.35E-11± 1.65E-12 | NS |
| *N*-docosahexaenoyl glycine | 8.7E-13± 1.15E-13 | 1.02E-12± 1.27E-13 | 8.61E-13± 1.26E-13 | 1.28E-12± 3.05E-13 | NS |
| ***N*-acyl taurine** | | | | | |
| *N*-palmitoyl taurine | 1.58E-10± 1.54E-11 | 1.55E-10± 2.04E-11 | 1.37E-10± 9.47E-12 | 1.46E-10± 1.85E-11 | NS |
| *N*-stearoyl taurine | 6.34E-11± 7.14E-12 | 7.23E-11± 6.43E-12 | 7E-11± 8.21E-12 | 7.16E-11± 8.72E-12 | NS |
| *N*-oleoyl taurine | 1.73E-11± 1.97E-12 | 1.85E-11± 2.48E-12 | 1.78E-11± 1.9E-12 | 1.63E-11± 2.27E-12 | NS |
| *N*-arachidonoyl taurine | 3.8E-11± 4.7E-12 | 3.75E-11± 5.66E-12 | 4.43E-11± 3.95E-12 | 3.83E-11± 4.49E-12 | NS |
| **2-acyl glycerol** | | | | | |
| 2-palmitoyl glycerol | 6.56E-09± 1.06E-09 | 7.76E-09± 1.15E-09 | 4.65E-09± 7.35E-10 | 5.54E-09± 8.32E-10 | NS |
| 2-oleoyl glycerol | *1.4E-08± 2.24E-09* | 1.26E-08± 2.26E-09 | *7.33E-09± 1.26E-09* | 8.59E-09± 1.55E-09 | *0.029* |
| 2-linoleoyl glycerol | *9.55E-11± 6.97E-12* | **9.31E-11± 8.44E-12** | *5.05E-11± 4.97E-12* | **4.95E-11± 4.94E-12** | *<0.001* **<0.001** |
| 2-arachidonoyl glycerol | *5.52E-10± 7.7E-11* | 4.39E-10± 7.76E-11 | *3.19E-10± 6.8E-11* | 3.3E-10± 5.33E-11 | *0.034* |

| | | | | | |
|---|---|---|---|---|---|
| **Free Fatty Acids** | | | | | |
| Oleic acid | 4.8E-09± 4.75E-10 | 5.52E-09± 5.33E-10 | 4.5E-09± 7.6E-10 | 5.13E-09± 5.51E-10 | NS |
| Linoleic acid | 5.04E-10± 2.96E-11 | 5.63E-10± 4.11E-11 | 4.22E-10± 3.64E-11 | 4.91E-10± 3.59E-11 | NS |
| Arachidonic acid | 3.13E-09± 1.12E-10 | 3.46E-09± 1.87E-10 | 3.49E-09± 3.05E-10 | 3.5E-09± 1.88E-10 | NS |
| **Prostaglandins** | | | | | |
| $PGE_2$ | 1.98E-11± 1.54E-12♂ | 2.88E-11± 1.82E-12♂ | 2.47E-11± 1.6E-12 | 2.53E-11± 2.34E-12 | 0.001♂ |
| $PGF_{2\alpha}$ | *2.71E-11± 1.16E-12*♂ | 3.75E-11± 9.46E-13♂ | *3.21E-11± 2.53E-12* | 3.33E-11± 1.41E-12 | <0.001♂ *0.027* |
| 6-keto$PGF_{1\alpha}$ | BAL | BAL | BAL | BAL | -- |
| **Sample Mass** | | | | | |
| Sample Mass | 0.02±7.32E-4 | 0.02±1.24E-3 | 0.02±6.29E-4 | 0.02±6.72E-4 | NS |

Data are moles per gram tissue and are shown as means ± SE. Values in light face have no significant difference among the groups. *Least Significant Difference corrected $p \leq 0.05$. *Italicized* values denote a significant sex effect for normally reared animals, whereas those in **bold** denote a significant sex effect for limited bedding animals; ♂=significant treatment effect for males, NS=Not Significant. BAL=Below Analytical Limits.
